# Architectural Groups of a Subtelomeric Gene Family Evolve Along Distinct Paths in *Candida albicans*

**DOI:** 10.1101/2022.08.15.504027

**Authors:** Matthew J. Dunn, Emily Simonton, Jason C. Slot, Matthew Z. Anderson

## Abstract

Subtelomeres are dynamic genomic regions shaped by elevated rates of recombination, mutation, and gene birth/death. These processes contribute to formation of lineage-specific gene family expansions that commonly occupy subtelomeres across eukaryotes. Investigating the evolution of subtelomeric gene families is complicated by the presence of repetitive DNA and high sequence similarity among gene family members that prevents accurate assembly from whole genome sequences. Here we investigated the evolution of the te<u>lo</u>mere-associated (*TLO*) gene family in *Candida albicans* using 189 complete coding sequences retrieved from 23 genetically diverse strains across the species. *TLO* genes conformed to the three major structural groups (α /β /γ) previously defined in the genome reference strain but significantly differed in the degree of within-group diversity and positional conservation. One group, *TLO*β, was always found at the same chromosome arm with strong sequence similarity among all strains. In contrast, diverse Tloα sequences have proliferated among chromosome arms. Tloγ genes formed seven primary clades that included each of the previously identified Tloγ genes from the genome reference strain with limited mobility among chromosome arms. Structural groups displayed regions of high conservation that resolved newly identified functional motifs, providing insight into potential regulatory mechanisms that distinguish groups. Thus, by resolving intra-species subtelomeric gene variation, it is possible to identify previously unknown gene family complexity that may underpin adaptive functional variation.

## INTRODUCTION

Gene duplication commonly gives rise to similar or identical paralogs through errors in DNA replication, sister chromatid exchange, or whole genome duplication (Tilley and Birshtein 1985; Kellis *et al*. 2004; Mehta and Haber 2014; Reams and Roth 2015). In most cases, one of the paralogs is inactivated by deleterious mutations, thereby restricting further evolutionary outcomes of paralogy (Cliften *et al*. 2006; Albalat and CaÑestro 2016). However, duplicate genes that remain functional may retain the ancestral function, split the ancestral function or interaction networks between paralogs, or evolve specialized or novel functions Hughes 1994; Wapinski *et al*. 2007; Des Marais and Rausher 2008; Innan and Kondrashov 2010). Retention of paralogs following repeated gene duplications can lead to the formation of a complex gene family whose members have the potential to diverge over evolutionary time.

Functional studies of gene family expansion have usually focused on paralog pairs in order to simplify experiments and inferences about selective pressures on genes following amplification (Kondrashov *et al*. 2002; Wagner 2002; Brunet *et al*. 2006; Cliften *et al*. 2006; Guan *et al*. 2007). Yeast species that have undergone whole genome duplication or regional mutations also make studies of many gene duplicates simultaneously convenient (Dietrich 2004; Kellis *et al*. 2004; Scannell *et al*. 2007a; Scannell *et al*. 2007b; Albertin and Marullo 2012). Studies of gene duplication demonstrated that functional outcomes are influenced by genomic context (Zhao and Boerwinkle 2002; Carreto *et al*. 2008; Zhu *et al*. 2014), gene dosage and protein complex formation (Aury *et al*. 2006; Veitia *et al*. 2008; Makino and McLysaght 2010), as well as by gene expression level (Aury *et al*. 2006; Conant and Wolfe 2006). However, the functional roles of individual paralogs from large gene families in fungi that expanded beyond a few copies remain largely unexplored, despite numerous developmentally and ecologically important gene family expansions (Brown *et al*. 2010; Floudas *et al*. 2012; Viragh *et al*. 2022).

Expanded gene families are often enriched in subtelomeric regions that are immediately adjacent to the telomeric repeats. Subtelomeres harbor a mixture of duplicated genes and repetitive sequences from fragmented mobile genetic elements (Corcoran *et al*. 1988; Riethman 2008; Kupiec 2014). In addition to copy number variation, subtelomeric genes are characterized by a rapid accumulation of mutations that can alter their expression, structure, or function (Brown *et al*. 2010). Frequent recombination, elevated mutation rates via acquisition of single nucleotide polymorphisms (SNPs) and insertions/deletions (indels), and the constant processes of gene duplication and disruption contribute to subtelomeres often being the most dynamic regions of the genome (Winzeler *et al*. 2003; Linardopoulou *et al*. 2005; Carreto *et al*. 2008; Kasuga *et al*. 2009; Anderson *et al*. 2015; Yue *et al*. 2017; Chen *et al*. 2018). Importantly, gene families that reside within the subtelomeres are often under selection from species-specific lifestyles (Mefford 2001; Dujon *et al*. 2004; Kyes *et al*. 2007; Linardopoulou *et al*. 2007; Brown *et al*. 2010; Chen *et al*. 2018; Otto *et al*. 2018). For example, the *MAL, MEL*, and *SUC* genes in *S. cerevisiae* allow cells to utilize different carbon sources (maltose, melibiose, and sucrose, respectively), and fluctuate in copy number depending on the available growth substrate (Brown *et al*. 2010; Wenger *et al*. 2011; Dunn *et al*. 2012).

The expansion of several gene families involved in virulence traits distinguishes *Candida albicans*, the most clinically relevant *Candida* species, from closely related yeasts. Expansion of the *ALS, SAP*, and *LIP* gene families in *C. albicans* increases the available repertoire of adhesins, proteases, and lipases, respectively, which contribute to host colonization and tissue destruction (Magee *et al*. 1993; Hube *et al*. 2000; Hoyer 2001). The most dramatic gene expansion occurred within the te<u>lo</u>mere-associated (*TLO*) gene family, which are present in fourteen copies in the *C. albicans* genome reference strain SC5314, two copies in the most closely-related *C. dubliniensis* species, and a single copy within all other *Candida* species (Butler *et al*. 2009; Jackson *et al*. 2009). All but one *TLO* gene are found in the subtelomeres of the eight *C. albicans* chromosomes where they often reside as the ultimate or penultimate gene. The fourteen *TLO* genes can be separated into three architectural groups (α, β, and γ) based on sequence variation that clusters towards the 3’ end of the gene (Van het Hoog *et al*. 2007; Anderson *et al*. 2012). *TLO* genes display high levels of sequence similarity. *TLO* paralogs have ∼97% nucleotide identity within a clade and 82% identity between clades when excluding indels (Van het Hoog *et al*. 2007; Anderson *et al*. 2012).

*TLO* genes encode a conserved N-terminal MED2 domain that facilitates their incorporation as stoichiometric components of the major transcriptional regulation complex Mediator (Zhang *et al*. 2012). Downstream of the MED2 domain is a gene-specific region of variable length followed by the 3’ portion of the gene that defines three *TLO* structural types (α /β /γ). *TLO*β *2* resides at the syntenic locus to *MED2* orthologs in other *Candida* species (Jackson *et al*. 2009) although *TLO*α group members appear to have given rise to *TLO*γ genes based on inferred mutational history (Anderson *et al*. 2012). The single *TLO*β group member contains two indels relative to *TLO*α group sequences, and *TLO*γ group members are defined by an LTR *rho* insertion that introduced a stop codon and truncated the coding sequence (Anderson *et al*. 2012). Recent diversification of these genes in *C. albicans* has resulted in variable *TLO* copy numbers among clinical isolates (Hirakawa *et al*. 2015), consistent with rapid gene loss/gain during *in vitro* passaging (Anderson *et al*. 2015).

Subtelomeric gene evolution in *C. albicans* has not been thoroughly explored at the individual sequence level because of the complexities in accurately resolving paralog gene structure and sequences from whole genome sequencing assemblies. Here we obtained complete sequences of the subtelomeric *TLO* genes in 23 well characterized strains. *TLO* sequences provide evidence for complex evolutionary histories among groups in this single gene family. Sequenced genes conformed to one of three previously defined structural groups (α, β, γ) with the exception of a small number of truncated open reading frames. We further identified strong conservation of a prion-like domain in a majority of Tloα and Tloβ sequences and two transmembrane regions in most Tloγ proteins. Surprisingly, phylogenetic analyses suggest that while Tloβ is monophyletic, Tloγ and truncated architectures may have emerged multiple times from the Tloα architecture. The degree of sequence divergence among groups varied significantly with high similarity among Tloβ sequences and high diversity among Tloα proteins. These evolutionary processes have resulted in diverse *TLO* repertoires among strains of *C. albicans* the functional consequences of which remain to be investigated.

## RESULTS

Short read sequencing of 23 *C. albicans* clinical isolates failed to accurately incorporate sequence variants into subtelomeric genes known to be present into the resulting assemblies (Hirakawa *et al*. 2015). These isolates capture much of the diversity present in *C. albicans* as they originate from different geographic regions, body sites of isolation, and a range of clades within the species (Butler *et al*. 2009; Hirakawa *et al*. 2015; Cuomo *et al*. 2019). To determine intra-species variation in *C. albicans* subtelomeric genes, we employed a chromosome-arm specific amplification and sequencing strategy that is capable of identifying any *TLO* gene present on a given chromosome arm through the use of centromeric chromosome arm-specific primers in combination with a primer that binds to a conserved sequence at the *TLO* start codon (Supplemental Figure S1). Resolved full length sequences facilitated characterization of gene architecture and mapping of structural and location data to a comprehensive gene phylogeny. This enabled the inference of trends in *TLO* molecular evolution and possible key events in the diversification of *TLO* genes across Candida.

### *Candida albicans* has a positionally and architecturally *diverse TLO* repertoire

*TLO*-specific amplification was performed for both subtelomeres of all eight *C. albicans* chromosomes in the 23 genetic backgrounds. Each of the resulting 299 amplicons were Sanger sequenced bidirectionally to produce 189 total full *TLO* gene sequences, representing between 4 and 14 products for each isolate (Figure 1A,B). Consistent with the genome reference strain SC5314, the right arm of chromosome 2 (Chr2R), Chr6R, and Chr7L did not produce any robust amplification products for any strain, indicating these chromosome arms do not encode *TLO* genes in *C. albicans*. All Tlo sequences contained an intact MED2 domain and were subsequently sorted into three *TLO* architectural groups based on similarity within a MAFFT alignment of the inferred protein sequences (Figure 1D). In total, 89 sequences conformed to the Tloα group gene architecture, 22 sequences to the Tloβ group, and 68 sequences to the Tloγ group. Conservation of specific Tlo sequences among related strains was evident in some cases but only in a minority of sequences (Figure 1C). The number of amplified *TLO* genes nor their relative representation among the three groups (α /β /γ) correlated with MLST clade designations (Supplemental Table 1). Additionally, nonsense mutations disrupted ten additional *TLO* sequences that are predicted to encode a complete MED2 domain but very little C-terminal peptide sequence and therefore did not conform to the α /β /γ group architecture.

**Figure 1.**
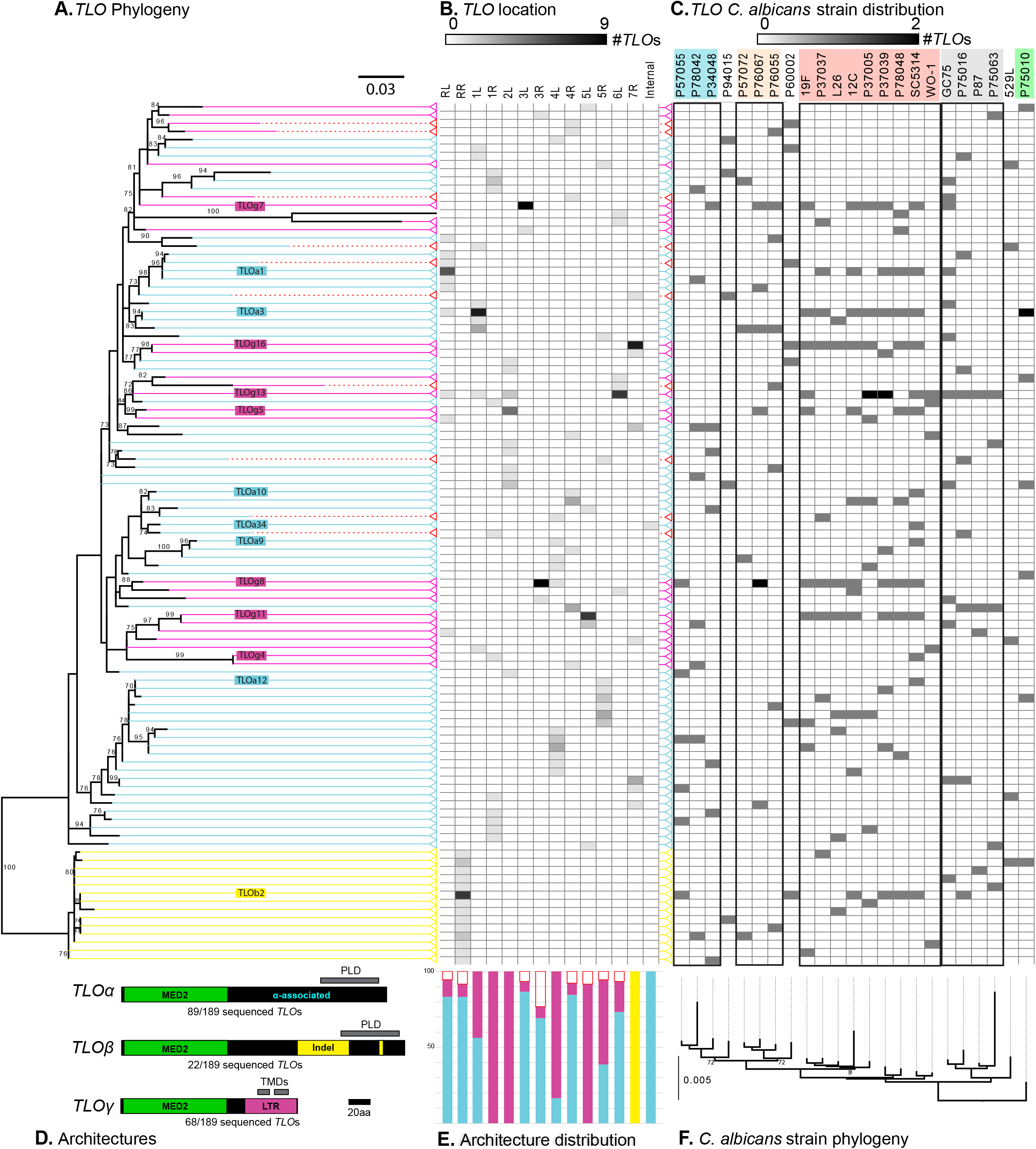
Tlo architectures across *Candida albicans* isolates. A maximum likelihood phylogeny of 189 translated *TLO* sequences was constructed under the JTTDCMut+G4 evolutionary model in IQTree (left). Identical sequences were collapsed into a single taxon prior to analysis. Support values are the percentage of 1000 IQTree ultrafast bootstrap (UFBoot) method. Terminals correspond to identical sequences with color indicating Tlo architectures (Tloα — solid cyan, Tloβ — solid yellow, Tloγ — solid pink, truncated – dashed red). The structure of each *TLO* architecture is depicted underneath the phylogeny. Unique sequence features separate 179 of 189 sequenced *TLO*s into three *TLO* architecture groups. Predicted prion-like domains (PLD) and transmembrane domains (TMD) are indicated with grey bars. The locations of *TLO* sequences are indicated as heat map shading for each chromosome arm (middle). The relative representation at each *TLO* position is indicated below (*TLO*α — solid cyan, *TLO*β — solid yellow, *TLO*γ — solid pink, truncated – red outline). Each Tlo sequence in the phylogeny is matched to the genetic background. A whole genome phylogeny is represented below. Clusters of genetically related strains are outlined in black boxes.

We obtained good representation of *TLO* sequences for most chromosome arms, which revealed a clear pattern of *TLO* gene representation. *TLO*β *2* genes were consistently recovered only from ChrRR. The other chromosome arms contained either only *TLO*γ sequences or a mix of *TLO*α and *TLO*γ genes. Extensive *TLO*α /γ group swapping was observed on Chr2L, Chr4R, and Chr7R, while the only intact loci on ChrRR, Chr3L, Chr3R, and Chr6L encoded *TLO*γ group members (Figure 1E).

Transcription factors containing prion-like domains (PLDs) can form phase-separated condensates in *C. albicans* that regulate cell identity (Frazer *et al*. 2020). We speculated similar molecular mechanisms may regulate transcriptional activators, including Tlo proteins. Indeed, 91 of 189 Tlos contained a predicted prion-like domain (PLD) (Figure 1 and Supplemental Table 2). PLDs were restricted to Tloα and Tloβ group proteins, although some Tloα and Tloβ sequences lacked a recognizable PLD (*e*.*g*., ChrRR in L26, ChrRR in P37005; Supplemental Figure S2).

Immunoprecipitation and mass spectrometry previously confirmed Tloα and Tloβ proteins associate with Mediator as predicted Med2 orthologs but failed to identify Mediator-bound Tloγ proteins (Zhang *et al*. 2012). Scanning the Tloγ sequences for unknown motifs uncovered two putative transmembrane domains (TMDs) in the C-terminal 50 amino acids (AA) in all but two Tloγ group members (Figure 1D). Specifically, the Tloγ sequence on Chr4R of P78042 contained only one predicted TM region, and none were predicted in Tloγ 4, a previously described Tloγ truncation on Chr1R of SC5314 (Anderson *et al*. 2012). The two predicted transmembrane helices are separated by a short 3 AA cytoplasmic loop that would place most of the Tloγ protein, including the Med2 domain, on the internal face of the mitochondrion.

### The *MED2* domain is highly conserved across *TLO* genes

The N-terminal MED2 domain defines *TLO* genes as Med2 homologs that incorporated as subunits of the larger Mediator complex (Yin and Wang 2014; Plaschka *et al*. 2016). HMMER search recovered the conserved MED2 domain in all 189 Tlo sequences and identified 90 homologous amino acid positions that are present in all sequences. Maximum likelihood phylogenetic analysis of the Tlo MED2 domain alignment did not recover distinct α /β /γ clades previously inferred using structural architecture (Supplemental Figure S2). Minimal branch lengths separated most Tlo MED2 sequences with the major exception of the *TLO*s on Chr6L in P37037 and P78048. These MED2 sequences are strongly separated from the rest of the phylogeny by amino acid variants that begin two thirds of the way through the MED2 domain (Chr6L in P37037; 71-90 AA, Chr6L in P78048; 62-90 AA).

### Monophyly is only strongly supported for the *TLO*β architecture

We built a MAFFT alignment of the homologous sequences among all *TLO* genes and further refined the output manually. Application of maximum likelihood to infer evolutionary relationships among architectures supported monophyly of Tloβ genes, whereas Tloα and Tloγ sequences were intermixed (Figure 1A). To test for monophyly of each Tlo architecture, constraint analyses (Table 1) were performed on each Tlo architecture independently and in all possible pairings. Constrained analyses that forced each individual architecture into a single node using the full dataset rejected the monophyly of the Tloα (Approximately unbiased (AU) test; p = 0.015) and truncated architectures (AU test; 0.021). Monophyly was not rejected for the *Tlo*β and Tloγ architectures using the full dataset, but monophyly of Tloγ was rejected in an alignment that excluded the truncated sequences (AU test; p = 2.09E-3). Assuming that *TLO*β is the ancestral architecture based on shared synteny between the ChrRR locus and chromosomes encoding *MED2* homologs in other *Candida* species (Jackson *et al*. 2009), these results suggest that *TLOα* genes originated once by duplication of *TLO*β, and *TLO*γ architectures arose from *TLOα*, possibly more than once. Truncated architectures arose from both *TLO*α and *TLO*γ sequences.

**Table 1.** Results of constraint analyses.

| Constraint | p-value* | LogL |
| --- | --- | --- |
| none (maximal likelihood tree) | 0.693 | -2983.985 |
| alpha monophyletic | **0.015** | -3054.839 |
| beta monophyletic | 0.583 | -2986.888 |
| gamma monophyletic | 0.134 | -3020.201 |
| beta gamma both monophyletic | 0.134 | -3020.201 |
| truncated monophyletic | **0.021** | -3030.799 |
| truncated free, architectures monophyletic | 0.086 | -3024.149 |
| group architectures each monophyletic | **0.004** | -3069.540 |
| *p-AU reported: p-value of approximately unbiased (AU) test (Shimodaira, 2002) |  |  |

### *TLO*β sequence architectures are conserved

Twenty-two sequences were identified as *TLO*β architectures based, in large part, on the presence of two defining indels towards the 3’ end of the gene when compared to other *TLO* architectures. Analysis of 189 Tloβ homologous positions revealed relatively little divergence among these sequences (Figure 2A). Only three positions encoded amino acid variants in the full alignment of all 22 Tloβ sequences (AA 117, 152, and 298 in the full alignment). Most variation in Tloβ sequences involved expansion or contraction of one of the defining indels rather than amino acid substitutions. Sequence variants in Tloβ genes clustered immediately after the MED2 domain in the first of two Tloβ -defining indels and corresponded to copy number variation of a tandem repeat that codes for a ‘TIDD/E’ amino acid sequence (Figure 2B). Eight Tloβ sequences contained one or two fewer ‘TIDD/E’ amino acid repeats compared to the SC5314 reference genome. The Tloβ sequence in isolate L26 contained a nonsense mutation at amino acid position 209, shortening this Tloβ protein by 63 amino acids but did not interrupt either the MED2 domain or the *TLO*β -specific indels.

**Figure 2.**
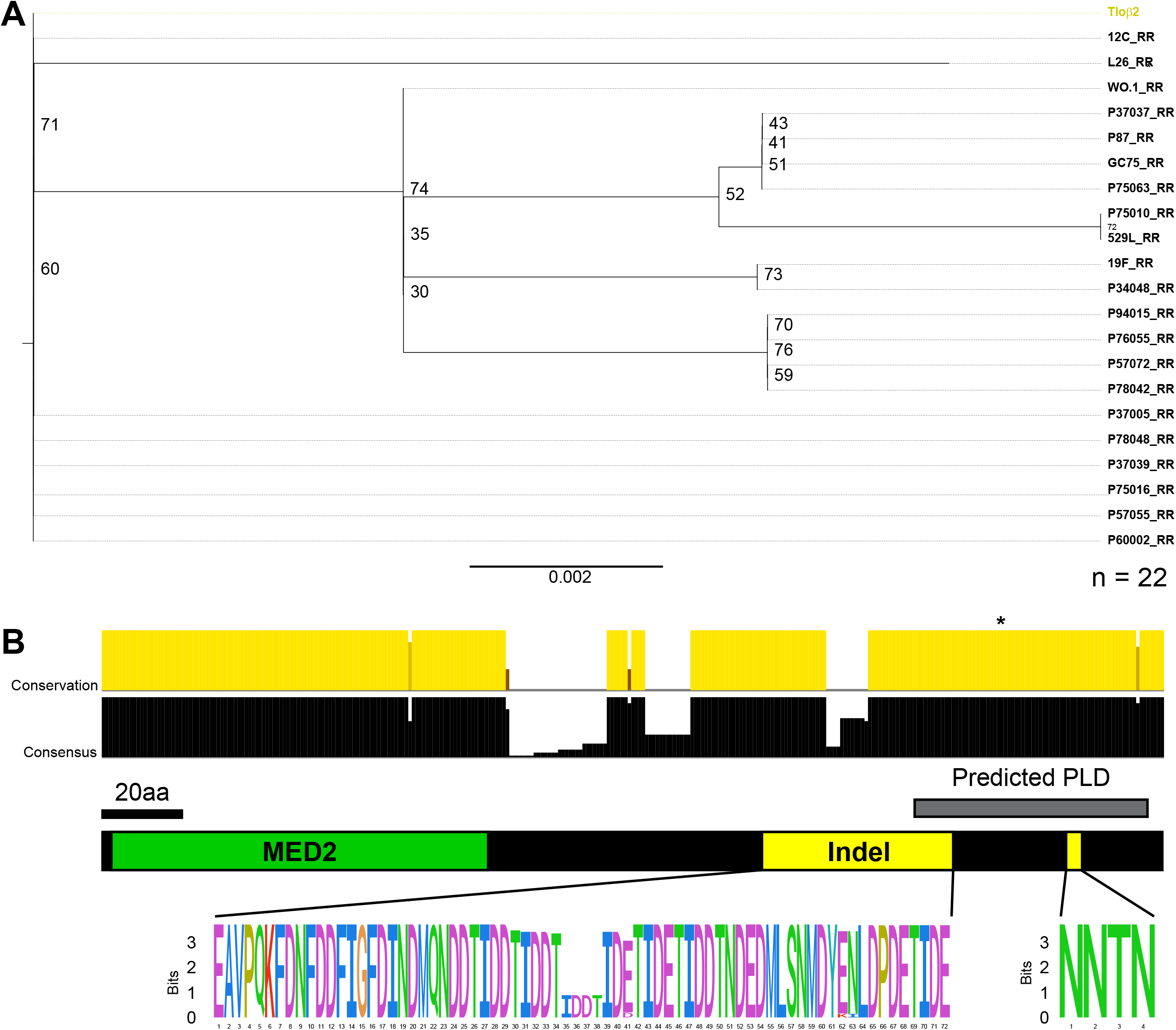
*TLO*β group genes display little variation. **A)** The 22 Tloβ group sequences were assessed for evolutionary relationships using maximum likelihood with JTT+G evolutionary models and 1000 bootstrap replicates. *TLO* sequences are reported as “Patient Isolate_Chromosome Arm.” Tlo sequence from SC5314 is colored. **B)** The canonical *TLO*β group structure is cartooned, where the two insertion events have been isolated as a sequence logo. Letters represent individual amino acids where the height signifies site representation. Amino acids within the sequence logo are colored using the default ClustalX coloring.

### *TLO*α genes are highly diversified

Radiating sequence diversity was present among *TLO*α members with most genes encoding a unique sequence relative to all other group members (Figure 3A). To identify other conserved functional domains in Tloα genes, we inferred the consensus alignment of all Tloα sequences. Two regions displayed high conservation in the alignment: the N-terminal MED2 domain and a second region covering the putative C-terminal PLD (Figure 3B). Truncation of four Tloα sequences by nonsense mutations occurs immediately downstream of the predicted PLD, but these genes still clustered with Tloα sequences.

**Figure 3.**
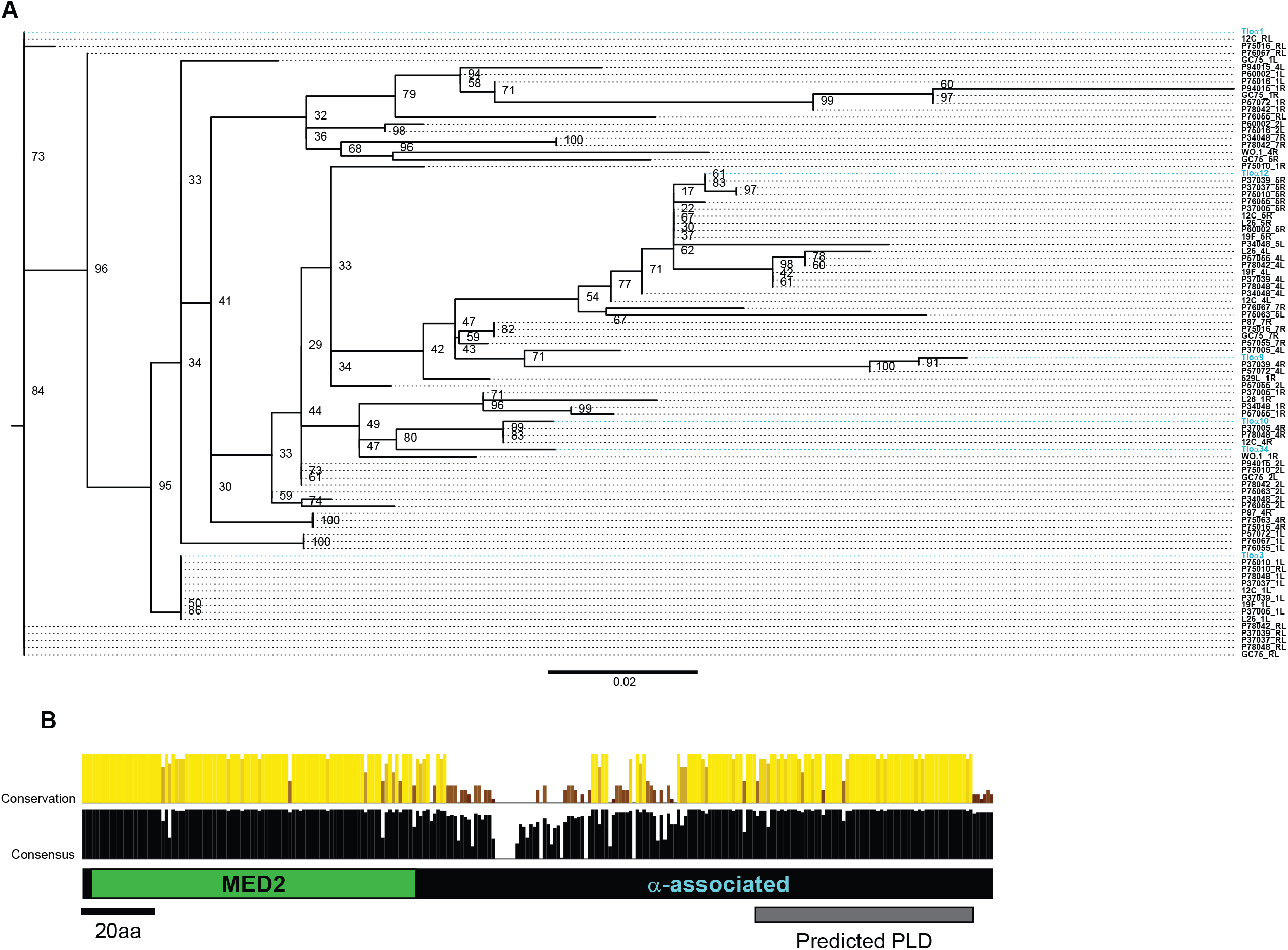
Expansive sequence variation among Tloα. **A)** Phylogenetic reconstruction as described in Figure 2 was conducted for the 89 Tloα group sequences and the resultant tree was visualized with FigTree. Tlo sequences are reported as “Patient Isolate_Chromosome Arm”. Tlo sequences from SC5314 are colored. **B**. The canonical Tloα group structure is cartooned with the Med2 and prion-like domain indicated. Conservation and consensus at each position are plotted at each position for all 89 Tloa sequences based on alignment using MAFFT. Bar height and strength of color signifies the strength at each position.

### *TLO*γ sequences cluster around gene-specific clades

The remaining 68 sequences in the dataset are truncated in an identical location by a single rho LTR 3’ insertion that defines the *TLO*γ architecture. The conserved sequence of the LTR insertion includes the two predicted TMDs. Altogether, 129 homologous sites were present in a Tloγ alignment that included 90 sites in the MED2 domain. The Tloγ -only phylogeny had moderate to strong bootstrap support at terminal nodes that contained highly similar sequences to single SC5314 Tloγ genes (Figure 4A). Each of these nodes typically contained genes from the same chromosome arm across strains indicating strong positional conservation within each Tloγ archetype in sharp contrast with the low positional conservation of the Tloα group. Removal of two truncated *TLO*γ sequences (*TLO*γ *4* and Chr4R in P78042) increased the total number of informative sites from 129 to 156 but did not significantly alter the phylogenetic relationships among the Tloγ sequences.

**Figure 4.**
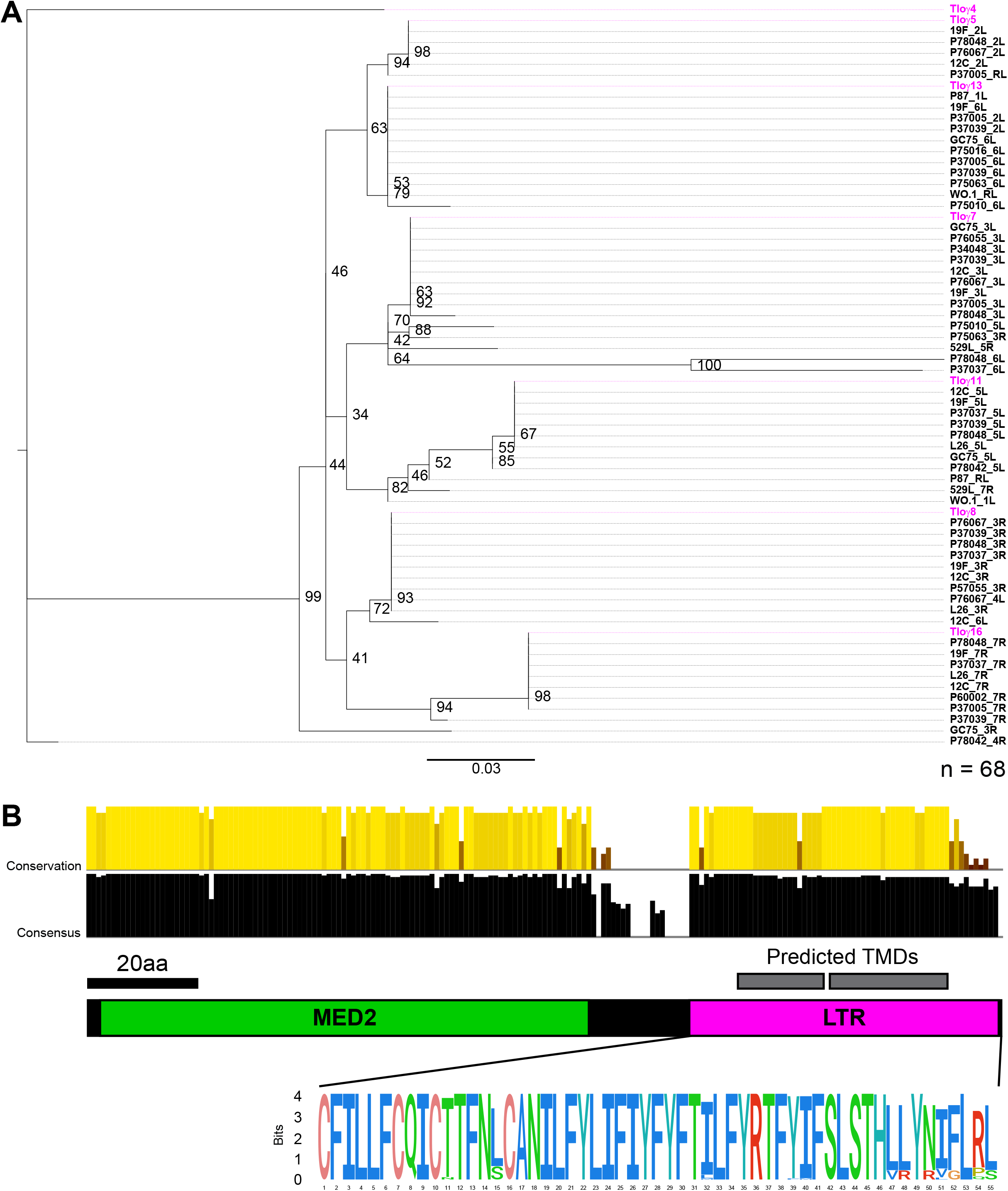
*TLO*γ group phylogenetic organization reveals gene clades. **A)** Phylogenetic reconstruction as described in Figure 2 was conducted for the 68 Tloγ group sequences and the resultant tree was visualized with FigTree. Tlo sequences are reported as “Patient Isolate_Chromosome Arm”. Tlo sequences from SC5314 are colored. **B)** The conserved Tloγ group structure has been cartooned, where the LTR has been isolated as a sequence logo. Letters represent individual amino acids where the height signifies site representation. Amino acids within the sequence logo are colored using the default ClustalX coloring.

Two-thirds of the Tloγ protein alignment (122/183 sites) is identical across sequences. Although the MED2 domain was expected to be conserved, the sequence within the Tloγ -defining *rho* LTR insertion was also surprisingly conserved (36/53 identical sites, Figure 4B). Nine of the 17 variant sites in the LTR insertion are due to Tloγ 16-like sequences that encode a distinct C-terminal nine amino acid peptide (VRYRVGLPS) with no notable similarity to other *C. albicans* genes.

*TLO*α -like sequences remain downstream of the rho LTR insertion that define the *TLO*γ architecture (Anderson *et al*. 2012). These sequences in SC5314 retain strong similarity to one another and the C-terminal end of Tloα proteins (Supplemental Figure S3A). Inclusion of these sequences in the Tlo phylogeny did not significantly alter the topology of the tree and interspersed placement of Tloα and Tloγ sequences (Figure S3B).

### Gene disrupting mutations do not show any clear patterns

Ten *TLO* genes contained ORF-disrupting mutations that significantly truncated the coding sequence. The *TLO* located on ChrRL in P60002 experienced a frameshift due to a single basepair insertion, while nonsense mutations disrupted all other *TLO* genes. Most of the genes contain a premature stop between 130 and 154 AA, shortly after the MED2 domain, where sequence conservation immediately declines at AA92 in the alignment (Supplemental Figure S3). It was possible to determine the architecture prior to truncation for each of these sequences, and this architecture is consistent with placement in the *TLO* phylogeny (Figure 1, Supplemental Table 3). The *TLO*s on Chr4R in P76055, P60002, and GC75 share a 34 amino acid C-terminal peptide with no similarity to any other *C. albicans* protein that we hypothesize is a result of Chr4R sequence divergence following truncation.

## DISCUSSION

Interrogation of expanded gene families often relies exclusively on variation present in the genome reference strain without considering additional intra-species sequence diversity. Genetic variation in the *C. albicans TLO* subtelomeric gene family among 23 clinical strains expands our understanding of the of evolutionary processes shaping the gene repertoire during gene family expansion. Expansion and differentiation of the *TLO* family into three groups has resulted in distinct sequence conservation outcomes among the three groups, ranging from strong conservation to diversification. Conserved segments of each *TLO* group alignment highlight previously overlooked functional domains that may contribute to functional diversification among paralogs. Together, this work reveals the balance between gene sequence diversification and novel functional motif conservation during subtelomeric gene family evolution that can confer unique attributes to gene subsets.

Little work has probed the sequence diversity in expanded gene families across strains of a single species and this is especially true for subtelomeric gene families. Although the importance of subtelomeric genes to unique organismal lifestyles and their accelerated evolution are well documented (Corcoran *et al*. 1988; Winzeler *et al*. 2003), investigation of genes in these regions has lagged because it is difficult to resolve their complex sequence structures and representation in repetitive subtelomeres across eukaryotic species (Riethman 2008; Young *et al*. 2020). We show here that traditional Sanger sequencing approaches are able to overcome limitations in Illumina short-read sequencing in accurately resolving these repetitive regions. Adoption of Nanopore and other high fidelity long-read sequencing will greatly facilitate investigations of subtelomeric gene family evolution to organismal adaptation as single reads can span full repetitive regions of the subtelomeres (Tyler *et al*. 2018; Zhou *et al*. 2019; Munoz *et al*. 2021).

Expansion of the *C. albicans TLO*s likely occurred through multiple independent events. Only the Tloβ architectural group is clearly monophyletic in *C. albicans* and may reflect its position as the ancestral *TLO* structure. The conserved position of *TLO*β *2* on ChrRR and the synteny of the position with *MED2* orthologs in other *Candida* species (Jackson *et al*. 2009) raises the hypothesis that it may maintain the ancestral function. In support of this, both *TLO*β *2* and the syntenic *Candida dubliniensis TLO1* contribute to filamentation (Haran *et al*. 2014; Uppuluri *et al*. 2018). The *TLO*α group appears then to have emerged multiple independent times in the *C. albicans* lineage via loss of the two indels that distinguish these architectural groups. If the TLO phylogeny is correct, it is most parsimonious to infer that the insertion of the rho LTR into the *TLO*α sequence to produce the *TLO*γ architecture occurred multiple times. However, the common insertional position into *TLO*α by the same retroelement type that resulted in *TLO*γ suggests either a single event or a region particularly vulnerable to disruption.

Most surprising from our analysis was that different *TLO* groups within the single expanded gene family have experienced disparate modes of sequence diversification. At one extreme, monophyletic Tloβ sequences have undergone very little sequence diversification despite our hypothesis that they represent the root of the *TLO* expansion. At the other extreme, Tloα proteins have undergone less constrained evolution and explore a wide swath of sequence space (Povolotskaya and Kondrashov 2010). This could reflect a longer amount of time for evolution to act on individual homologs or altered selective pressures on this gene architecture, which may no longer maintain the ancestral function. Lastly, the *TLO*γ sequences fall between these opposing extremes with clearly delineated clades that correspond to the *TLO*γ genes present in SC5314.

The most parsimonious explanation for the *TLO*γ architecture is that *TLO*γ paralogs have come under purifying selection following sequence diversification that occurred early in the diversification of *C. albicans* since many unrelated strains retain the same sequence in the same locus. That two paralogous genes, the ancestral *TLO*α and *TLO*β , have such different diversification patterns highlights the potential of gene family diversification to facilitate species adaptation. Organismal benefit may be derived from the alternative protein-protein interactions and transcriptional states conferred by incorporation of unique *TLO* structures and sequences into the transcriptional regulatory complex Mediator. Formation of alternate regulons by production of different Mediator types may operate as a bet hedging mechanism in the same cell or among cells in a population. Indeed, a vast excess of Tlo to Mediator in *C. albicans* (Zhang *et al*. 2012; Haran *et al*. 2014; Liu *et al*. 2016) could allow *TLO genes* to take unconstrained or additional evolutionary strategies independent of their orthologous role as Mediator components.

The molecular role of prion-like domain (PLD) in regulating availability of Mediator subunits has emerged as a common post-transcriptional regulatory mechanism (Zhu *et al*. 2015; Batlle *et al*. 2021). The PLD predicted in the majority of *TLO*α and *TLO*β paralogs contains a characteristic amyloid core required for phase transition of master regulators that define cell states in eukaryotic species (Hnisz *et al*. 2017). Recent work demonstrated similar mechanisms regulate *C. albicans* transcription factors (TFs) that govern transition between the white and opaque cell states (Frazer *et al*. 2020). Given that *TLO*α and *TLO*β sequences function interchangeably as the Med2 subunit of Mediator (Zhang *et al*. 2012; Zhang *et al*. 2013), it is tempting to speculate that Tlos form liquid-like droplets to sequester excess Tlo from Mediator or with Mediator itself. Sequestration of Tlo may alter general transcriptional activity of Mediator or change the patterns of RNA Polymerase II (PolII) recruitment to promoters by increasing association of Mediator with a subset of available Tlos (Haran *et al*. 2014).

Alignment of the Tloγ revealed the strong conservation over the LTR insertion that defines this *TLO* group. Conservation of this insertion indicated an embedded functional domain that led to the identification of transmembrane domains that may anchor Tloγ proteins in the outer mitochondrial membrane or internally in cristae. How their putative membrane association contributes to their ascribed function in Mediator is unclear since this complex is expected to require free diffusion and may suggest a mitochondrial function completely independent of its canonical role in Mediator (Mamouei *et al*. 2021).

Truncation of ten genes by nonsense mutations appears to have resulted in MED2 domains with little downstream sequence. *MED2* homologs in ascomycetes tend to contain an N-terminal MED2 domain followed by an extended C-terminal tail. Yet, the Med29 metazoan counterpart to fungal Med2 lacks the C-terminal extension in its role in the Mediator tail (Rengachari *et al*. 2021). Retention of the MED2 domain should allow association with Mediator, but how this affects recruitment of PolII and gene expression when lacking the C-terminal end to interact with transcription factors is unclear.

This investigation reinforced previous work showing frequent “movement” of *TLO* paralogs between chromosome arms (Anderson *et al*. 2015). Interestingly, the chromosome arms containing *TLO*α group sequences also always contain *TLO*γ sequences in other isolates, suggesting that the more recently emerged *TLO*γ genes may be more flexible in occupying various chromosomal position compared to *TLO*α genes. This is consistent with the unidirectional invasion and replacement of *TLO*α by *TLO*γ genes during passaging experiments with SC5314 (Anderson *et al*. 2015). While an eventual complete replacement of *TLO*α group members by *TLO*γ genes may be expected in this framework, a divergent function of *TLO*γ genes from *TLO*α paralogs may restrict their abundance among the *TLO* repertoire. Lastly, the placement of *TLO*β is consistently on ChrRR in each sequenced isolate may have resulted from the absence of a telomere recombination element (TRE) adjacent to *TLO*β *2*, previously noted in SC5314 (Freire-BenÉitez *et al*. 2016). Disruption of a TRE reduced rates of loss of heterozygosity on single chromosome arms and may similarly reduce interchromosomal recombination and gene “movement” to other chromosome arms when absent.

Altogether, this work demonstrates that subtelomeric gene family diversity is likely significantly underrepresented when using a single genome reference strain for eukaryotic species. As a result, current perspectives of genome evolution in functional subtelomeric sequences may be incomplete or skewed based on the limited data available in a single isolate. As seen for *C. albicans TLO* genes, expansion to include a strain collection revealed sequence diversification and the evolutionary histories of individual or groups of genes that were otherwise hidden.

## MATERIALS AND METHODS

### Amplification and Sequencing of Individual TLO paralogs

Strains of *Candida* used in this study are listed in Supplemental Table 4. Overnight cultures of each *C. albicans* strain were grown overnight in 3 mL of liquid YPD medium on a rotary drum at 30°C. DNA was purified from these cultures using the MasterPure™ Yeast DNA Purification Kit (Epicenter/Lucigen). Purified DNA was used to amplify each *TLO* gene using a centromeric chromosome arm specific primer (ALO36-48 and ALO60) paired with a conserved *TLO* start site primer (ALO35) (Supplemental Figure S1). Primer sequences are listed in Supplemental Table 5. Amplified products were examined on agarose gels by electrophoresis to confirm single, clean amplicons. Single-product amplification reactions were then purified using a magnetic bead approach (Berensmeier 2006). Purified DNA was sent for Sanger sequencing at the Genomics Shared Resource within the Ohio State University Comprehensive Cancer Center.

Sequencing was performed with primers oriented towards the *TLO* start site (ALO49-59 and ALO61) to sequence the amplified product (Supplemental Figure S1). Chr4R was sequenced using the *TLO* start site primer. A minimum of two independent sequencing reactions were performed for each *TLO* amplicon. The conserved start site primer was used to resolve any ambiguous sequence near the start codon.

### Phylogenetic Reconstructions

Consensus *TLO* nucleotide sequences were translated into amino acid sequences using the CUG fungal translation table (alternative yeast nuclear code - translation table 12). Sorted alignments were built from the 189 consensus Tlo sequences using MAFFT v.7 (Katoh and Standley 2013). Sequences were divided by group architecture (α, β, γ); where *TLO*β -like sequences contained one large and one small insertion, *TLO*γ -like sequences were interrupted by an LTR, and *TLO*α -like sequences contained neither of these events. Domain extraction was conducted for all full length sequences and also in isolation of the MED2 domain using HMMER v3.3.2 (Eddy 2011). ModelFinder (Kalyaanamoorthy *et al*. 2017) within IQ-TREE (Nguyen *et al*. 2015) was used for evolutionary model testing. Maximum likelihood phylogenies were run with 1000 bootstrap replicates within IQ-TREE using the ultrafast bootstrap (UFBoot) (Hoang *et al*. 2018) method under the best evolutionary model. Bootstrapped trees were then exported as Newick trees for visualization.

### Constraint Analyses

Monophyletic node architectures (Supplemental Table S6) were constructed in Mesquite v3.70 (http://www.mesquiteproject.org). Constrained topologies generated in IQ-TREE were compared by the Direct Computation with Mutabilities revised JTT model (JTTDCmut) with a gamma of four categories (Kosiol and Goldman 2005; Nguyen *et al*. 2015). Statistical analysis of these constraints is reported in Table 1.

### Prion-like Domain Identification

PrionW (Zambrano *et al*. 2015) was used to predict prion-like domains (PLDs) based on an amyloid core and predicted pWALTZ score. PLAAC analysis (Lancaster *et al*. 2014) was used to identify the strength to which PLD calls conformed to the canonical yeast PLD architecture based on hidden markov modelling (Supplemental Figure S4).

### Identification of Predicted Transmembrane Domains in Tlog Group Members

The Protein Homology/analogY Recognition Engine V 2.0 (Phyre2) web portal (Kelley *et al*. 2015) was used on “Expert Mode” to generate structural predictions for the 68 Tloγ group members from *Candida albicans*. A FASTA file containing the 68 Tloγ protein amino acid sequences was edited to remove any gaps and non-letter characters before it was submitted to the “Batch Processing” portal of Phyre2. Default parameters were used when applicable. Domain regions were then mapped back to the Tloγ group member MAFFT alignment for visualization.

### Data Visualization

Data visualization was conducted using R version 3.6.3. Bar charts were generated using Microsoft Excel. Newick format trees were visualized using FigTree v1.4.4 (http://tree.bio.ed.ac.uk/software/figtree/). Sequence conservation and consensus were visualized using Jalview (Waterhouse *et al*. 2009). Amino acid sequence logos were visualized using the R package ‘ggseqlogo’ (Wagih 2017) and colored using the default ClustalX coloring scheme (Thompson *et al*. 1994).

### Data Availability

All sequences used as part of this manuscript are included as a Supplemental File and available online at https://drive.google.com/file/d/1_UJ_GlqLEVdXhBwcjSBUsNHqPDL8ktrd/view?usp=sharing.

## ACKNOWLEDGEMENTS

We would like to thank the Genomics Shared Resource within the Ohio State University Comprehensive Cancer Center for their support with these sequencing efforts. Additionally, we would like to thank the Anderson and Rappleye lab members for their helpful feedback on data visualization and support through this project.

## AUTHOR CONTRIBUTIONS

Conceptualization: MJD, JCS, and MZA. Data curation: MJD. Formal analysis: MJD, JCS, and MZA. Funding acquisition: MJD and MZA. Investigation: MJD. Methodology: MJD, JCS, and MZA. Project administration: MZA. Resources: MZA. Software: MJD, JCS, and MZA. Supervision: JCS and MZA. Validation: MJD, JCS, and MZA. Visualization: MJD. Writing - original draft: MJD. Writing - review & editing: JCS and MZA.

## FIGURE LEGENDS

**Supplemental Figure 1. *TLO* sequencing strategy**. The strategy used to amplify *TLO* gene sequences from specific chromosome arms has been cartooned where relative locations of each of the primer sets used are specified. Complete primer list can be found in Supplemental Table 2.

**Supplemental Figure 2. The *TLO* MED2 domain is evolving independent of the C-terminal sequence architecture. A)** Phylogenetic reconstruction of the HMMER extracted MED2 domain was conducted on the 189 sequenced *TLO*s using 90 homologous sites. Maximum likelihood phylogenies were constructed using JTT+G evolutionary models with 1000 bootstrap replicates. Coloration was assigned by *TLO* group, where cyan represents *TLO*α members, yellow represents *TLO*β members, magenta represents *TLO*γ members, and red represents Tlo sequences outside of these three groups. Grey coloration indicates the presence of a prion-like domain (PLD). *TLO* sequences are reported as “Patient Isolate_Chromosome Arm.” Bootstrap values greater than 0.5 are shown at their respective nodes.

**Supplemental Figure 3. Conserved Tloα sequences lie downstream of *TLO*γ genes. A**. The region downstream of the architecture-defining *TLO*γ LTR insertion was translated for SC5314 and aligned with Tloα sequences using MAFFT. The conservation, quality, consensus, and occupancy at each site is given below. The intensity of blue shading indicates the similarity to the consensus residue at that position. **B**. The evolutionary relationship of all 189 Tlo sequences and the reconstructed “Tloγ “ genes prior to LTR disruption were assessed using maximum likelihood with JTT+G evolutionary models and 1000 bootstrap replicates. *TLO* sequences are reported as “Patient Isolate_Chromosome Arm.”

**Supplemental Figure 4. Truncated Tlo sequences are disrupted at the C-terminus**. The ten truncated Tlo sequences were aligned using MAFFT and the conservation and consensus are indicated in each line where bar height and strength of color signifiy the strength at each position.

**Supplemental Figure 5. Graphical representation of putative prion-like domains in Tlo proteins**. The results of PLAAC analysis for Tloα 12 and Tloβ 2 from the SC5314 genetic background are displayed.

**Supplemental Table 1. *TLO* sequence architectures are not defined by their MLST genotype group**.

**Supplemental Table 2. *TLO*α and *TLO*β group architectures contain PLD signatures in the 3’ architecture-specific region**.

**Supplemental Table 3. Assignment of sequences downstream of ORF-disrupting mutations**.

**Supplemental Table 4. Strains used in this study**.

**Supplemental Table 5. Oligonucleotides used in this study**.

**Supplemental Table 6. Constraint analysis by architecture and chromosomal position**.

## REFERENCES

Albalat, R., and C. Cañestro, 2016 Evolution by gene loss. Nature Reviews Genetics 17: 379–391.

Albertin, W., and P. Marullo, 2012 Polyploidy in fungi: evolution after whole-genome duplication. Proceedings of the Royal Society B: Biological Sciences 279: 2497–2509.

Anderson, M. Z., J. A. Baller, K. Dulmage, L. Wigen and J. Berman, 2012 The three clades of the telomere-associated Tlo gene family of Candida albicans have different splicing, localization, and expression features. Eukaryotic Cell 11: 1268–1275.

Anderson, M. Z., L. J. Wigen, L. S. Burrack and J. Berman, 2015 Real-Time Evolution of a Subtelomeric Gene Family in Candida albicans. Genetics 200: 907–919.

Aury, J.-M., O. Jaillon, L. Duret, B. Noel, C. Jubin et al., 2006 Global trends of whole-genome duplications revealed by the ciliate Paramecium tetraurelia. Nature 444: 171–178.

Batlle, C., I. Calvo, V. Iglesias, J. L. C M. Gil-Garcia et al., 2021 MED15 prion-like domain forms a coiled-coil responsible for its amyloid conversion and propagation. Commun Biol 4: 414.

Berensmeier, S., 2006 Magnetic particles for the separation and purification of nucleic acids. Applied Microbiology and Biotechnology 73: 495–504.

Brown, C. A., A. W. Murray and K. J. Verstrepen, 2010 Rapid expansion and functional divergence of subtelomeric gene families in yeasts. Current biology : CB 20: 895–903.

Brunet, F. d. r. G., H. R. Crollius, M. Paris, J.-M. Aury, P. Gibert et al., 2006 Gene Loss and Evolutionary Rates Following Whole-Genome Duplication in Teleost Fishes. Molecular Biology and Evolution 23: 1808–1816.

Butler, G., M. D. Rasmussen, M. F. Lin, M. A. S. Santos, S. Sakthikumar et al., 2009 Evolution of pathogenicity and sexual reproduction in eight Candida genomes. Nature 459: 657–662.

Carreto, L., M. F. Eiriz, A. C. Gomes, P. M. Pereira, D. Schuller et al., 2008 Comparative genomics of wild type yeast strains unveils important genome diversity. BMC genomics 9: 524.

Chen, N. W. G., V. Thareau, T. Ribeiro, G. Magdelenat, T. Ashfield et al., 2018 Common Bean Subtelomeres Are Hot Spots of Recombination and Favor Resistance Gene Evolution. Front Plant Sci 9: 1185.

Cliften, P. F., R. S. Fulton, R. K. Wilson and M. Johnston, 2006 After the Duplication: Gene Loss and Adaptation in Saccharomyces Genomes. Genetics 172: 863–872.

Conant, G. C., and K. H. Wolfe, 2006 Functional Partitioning of Yeast Co-Expression Networks after Genome Duplication. PLoS Biology 4: e109.

Corcoran, L. M., J. K. Thompson, D. Walliker and D. J. Kemp, 1988 Homologous recombination within subtelomeric repeat sequences generates chromosome size polymorphisms in P. falciparum. Cell 53: 807–813.

Cuomo, C. A., S. Fanning, S. Gujja, Q. Zeng, J. R. Naglik et al., 2019 Genome Sequence for Candida albicans Clinical Oral Isolate 529L Microbiology Resource Announcements 8: 20–21.

Des Marais, D. L., and M. D. Rausher, 2008 Escape from adaptive conflict after duplication in an anthocyanin pathway gene. Nature 454: 762–765.

Dietrich, F. S., 2004 The Ashbya gossypii Genome as a Tool for Mapping the Ancient Saccharomyces cerevisiae Genome. Science 304: 304–307.

Dujon, B., D. Sherman, G. Fischer, P. Durrens, S. Casaregola et al., 2004 Genome evolution in yeasts. Nature 430: 35–44.

Dunn, B., C. Richter, D. J. Kvitek, T. Pugh and G. Sherlock, 2012 Analysis of the Saccharomyces cerevisiae pan-genome reveals a pool of copy number variants distributed in diverse yeast strains from differing industrial environments. Genome Research 22: 908–924.

Eddy, S. R., 2011 Accelerated Profile HMM Searches. PLoS Computational Biology 7: e1002195.

Floudas, D., M. Binder, R. Riley, K. Barry, R. A. Blanchette et al., 2012 The Paleozoic origin of enzymatic lignin decomposition reconstructed from 31 fungal genomes. Science 336: 1715–1719.

Frazer, C., M. I. Staples, Y. Kim, M. Hirakawa, M. A. Dowell et al., 2020 Epigenetic cell fate in Candida albicans is controlled by transcription factor condensates acting at super-enhancer-like elements. Nature Microbiology 5: 1374–1389.

Freire-Benéitez, V., S. Gourlay, J. Berman and A. Buscaino, 2016 Sir2 regulates stability of repetitive domains differentially in the human fungal pathogen Candida albicans. Nucleic Acids Research 44: gkw594.

Guan, Y., M. J. Dunham and O. G. Troyanskaya, 2007 Functional Analysis of Gene Duplications in Saccharomyces cerevisiae. Genetics 175: 933–943.

Haran, J., H. Boyle, K. Hokamp, T. Yeomans, Z. Liu et al., 2014 Telomeric ORFs (TLOs) in Candida spp.

Encode Mediator Subunits That Regulate Distinct Virulence Traits. PLoS Genetics 10.

Hirakawa, M. P., D. A. Martinez, S. Sakthikumar, M. Z. Anderson, A. Berlin et al., 2015 Genetic and phenotypic intra-species variation in <i>Candida albicans</i>. Genome Research 25: 413–425.

Hnisz, D., K. Shrinivas, R. A. Young, A. K. Chakraborty and P. A. Sharp, 2017 A Phase Separation Model for Transcriptional Control. Cell 169: 13–23.

Hoang, D. T., O. Chernomor, A. Von Haeseler, B. Q. Minh and L. S. Vinh, 2018 UFBoot2: Improving the ultrafast bootstrap approximation. Molecular Biology and Evolution 35: 518–522.

Hoyer, L. L., 2001 The ALS gene family of Candida albicans. Trends in Microbiology 9: 176–180.

Hube, B., F. Stehr, M. Bossenz, A. Mazur, M. Kretschmar et al., 2000 Secreted lipases of Candida albicans : cloning, characterisation and expression analysis of a new gene family with at least ten members. Archives of Microbiology 174: 362–374.

Hughes, A. L., 1994 The evolution of functionally novel proteins after gene duplication. Proceedings of the Royal Society B: Biological Sciences 256: 119–124.

Innan, H., and F. Kondrashov, 2010 The evolution of gene duplications: classifying and distinguishing between models. Nature Reviews Genetics 11: 97–108.

Jackson, A. P., J. A. Gamble, T. Yeomans, G. P. Moran, D. Saunders et al., 2009 Comparative genomics of the fungal pathogens Candida dubliniensis and Candida albicans. Genome Research 19: 2231–2244.

Kalyaanamoorthy, S., B. Q. Minh, T. K. F. Wong, A. Von Haeseler and L. S. Jermiin, 2017 ModelFinder: Fast model selection for accurate phylogenetic estimates. Nature Methods 14: 587–589.

Kasuga, T., G. Mannhaupt and N. L. Glass, 2009 Relationship between Phylogenetic Distribution and Genomic Features in Neurospora crassa. PLoS ONE 4: e5286.

Katoh, K., and D. M. Standley, 2013 MAFFT Multiple Sequence Alignment Software Version 7: Improvements in Performance and Usability. Molecular Biology and Evolution 30: 772–780.

Kelley, L. A., S. Mezulis, C. M. Yates, M. N. Wass and M. J. E. Sternberg, 2015 The Phyre2 web portal for protein modeling, prediction and analysis. Nature Protocols 10: 845–858.

Kellis, M., B. W. Birren and E. S. Lander, 2004 Proof and evolutionary analysis of ancient genome duplication in the yeast Saccharomyces cerevisiae. Nature 428: 617–624.

Kondrashov, F. A., I. B. Rogozin, Y. I. Wolf and E. V. Koonin, 2002 Selection in the evolution of gene duplications. Genome biology 3: RESEARCH0008.

Kosiol, C., and N. Goldman, 2005 Different versions of the Dayhoff rate matrix. Mol Biol Evol 22: 193–199.

Kupiec, M., 2014 Biology of telomeres: lessons from budding yeast. FEMS Microbiology Reviews 38: 144–171.

Kyes, S. A., S. M. Kraemer and J. D. Smith, 2007 Antigenic Variation in Plasmodium falciparum: Gene Organization and Regulation of the var Multigene Family. Eukaryotic Cell 6: 1511–1520.

Lancaster, A. K., A. Nutter-Upham, S. Lindquist and O. D. King, 2014 PLAAC: a web and command-line application to identify proteins with prion-like amino acid composition. Bioinformatics 30: 2501–2502.

Linardopoulou, E. V., S. S. Parghi, C. Friedman, G. E. Osborn, S. M. Parkhurst et al., 2007 Human subtelomeric WASH genes encode a new subclass of the WASP family. PLoS Genet 3: e237.

Linardopoulou, E. V., E. M. Williams, Y. Fan, C. Friedman, J. M. Young et al., 2005 Human subtelomeres are hot spots of interchromosomal recombination and segmental duplication. Nature 437: 94–100.

Liu, Z., G. P. Moran, D. J. Sullivan, D. M. MacCallum and L. C. Myers, 2016 Amplification of TLO Mediator Subunit Genes Facilitate Filamentous Growth in Candida Spp. PLOS Genetics 12: e1006373.

Magee, B. B., B. Hube, R. J. Wright, P. J. Sullivan and P. T. Magee, 1993 The genes encoding the secreted aspartyl proteinases of Candida albicans constitute a family with at least three members. Infection and Immunity 61: 3240–3243.

Makino, T., and A. McLysaght, 2010 Ohnologs in the human genome are dosage balanced and frequently associated with disease. Proceedings of the National Academy of Sciences 107: 9270–9274.

Mamouei, Z., S. Singh, B. Lemire, Y. Gu, A. Alqarihi et al., 2021 An evolutionarily diverged mitochondrial protein controls biofilm growth and virulence in Candida albicans. PLoS Biology 19: 1–27.

Mefford, H. C., 2001 Comparative sequencing of a multicopy subtelomeric region containing olfactory receptor genes reveals multiple interactions between non-homologous chromosomes. Human Molecular Genetics 10: 2363–2372.

Mehta, A., and J. E. Haber, 2014 Sources of DNA Double-Strand Breaks and Models of Recombinational DNA Repair. Cold Spring Harbor Perspectives in Biology 6: a016428–a016428.

Munoz, J. F., R. M. Welsh, T. Shea, D. Batra, L. Gade et al., 2021 Clade-specific chromosomal rearrangements and loss of subtelomeric adhesins in Candida auris. Genetics 218.

Nguyen, L. T., H. A. Schmidt, A. Von Haeseler and B. Q. Minh, 2015 IQ-TREE: A fast and effective stochastic algorithm for estimating maximum-likelihood phylogenies. Molecular Biology and Evolution 32: 268–274.

Otto, T. D., U. Bohme, M. Sanders, A. Reid, E. I. Bruske et al., 2018 Long read assemblies of geographically dispersed Plasmodium falciparum isolates reveal highly structured subtelomeres. Wellcome Open Res 3: 52.

Plaschka, C., K. Nozawa and P. Cramer, 2016 Mediator Architecture and RNA Polymerase II Interaction. Journal of Molecular Biology 428: 2569–2574.

Povolotskaya, I. S., and F. A. Kondrashov, 2010 Sequence space and the ongoing expansion of the protein universe. Nature 465: 922–926.

Reams, A. B., and J. R. Roth, 2015 Mechanisms of Gene Duplication and Amplification. Cold Spring Harbor Perspectives in Biology 7: a016592.

Rengachari, S., S. Schilbach, S. Aibara, C. Dienemann and P. Cramer, 2021 Structure of the human Mediator-RNA polymerase II pre-initiation complex. Nature 594: 129–133.

Riethman, H., 2008 Human subtelomeric copy number variations. Cytogenetic and Genome Research 123: 244–252.

Scannell, D. R., G. Butler and K. H. Wolfe, 2007a Yeast genome evolution—the origin of the species. Yeast 24: 929–942.

Scannell, D. R., A. C. Frank, G. C. Conant, K. P. Byrne, M. Woolfit et al., 2007b Independent sorting-out of thousands of duplicated gene pairs in two yeast species descended from a whole-genome duplication. Proceedings of the National Academy of Sciences 104: 8397–8402.

Thompson, J. D., D. G. Higgins and T. J. Gibson, 1994 CLUSTAL W: improving the sensitivity of progressive multiple sequence alignment through sequence weighting, position-specific gap penalties and weight matrix choice. Nucleic Acids Research 22: 4673–4680.

Tilley, S. A., and B. K. Birshtein, 1985 Unequal sister chromatid exchange. A mechanism affecting Ig gene arrangement and expression. Journal of Experimental Medicine 162: 675–694.

Tyler, A. D., L. Mataseje, C. J. Urfano, L. Schmidt, K. S. Antonation et al., 2018 Evaluation of Oxford Nanopore’s MinION Sequencing Device for Microbial Whole Genome Sequencing Applications. Scientific Reports 8: 10931.

Uppuluri, P., M. Acosta Zaldívar, M. Z. Anderson, M. J. Dunn, J. Berman et al., 2018 Candida albicans Dispersed Cells Are Developmentally Distinct from Biofilm and Planktonic Cells. mBio 9: 1–16.

Van het Hoog, M., T. J. Rast, M. Martchenko, S. Grindle, D. Dignard et al., 2007 Assembly of the Candida albicans genome into sixteen supercontigs aligned on the eight chromosomes. Genome Biology 8.

Veitia, R. A., S. Bottani and J. A. Birchler, 2008 Cellular reactions to gene dosage imbalance: genomic, transcriptomic and proteomic effects. Trends in Genetics 24: 390–397.

Viragh, M., Z. Merenyi, A. Csernetics, C. Foldi, N. Sahu et al., 2022 Evolutionary Morphogenesis of Sexual Fruiting Bodies in Basidiomycota: Toward a New Evo-Devo Synthesis. Microbiol Mol Biol Rev 86: e0001921.

Wagih, O., 2017 ggseqlogo: a versatile R package for drawing sequence logos. Bioinformatics 33: 3645–3647.

Wagner, A., 2002 Selection and gene duplication: a view from the genome. Genome biology 3: reviews1012.

Wapinski, I., A. Pfeffer, N. Friedman and A. Regev, 2007 Natural history and evolutionary principles of gene duplication in fungi. Nature 449: 54–61.

Waterhouse, A. M., J. B. Procter, D. M. A. Martin, M. Clamp and G. J. Barton, 2009 Jalview Version 2--a multiple sequence alignment editor and analysis workbench. Bioinformatics 25: 1189–1191.

Wenger, J. W., J. Piotrowski, S. Nagarajan, K. Chiotti, G. Sherlock et al., 2011 Hunger Artists: Yeast Adapted to Carbon Limitation Show Trade-Offs under Carbon Sufficiency. PLoS Genetics 7: e1002202.

Winzeler, E. A., C. I. Castillo-Davis, G. Oshiro, D. Liang, D. R. Richards et al., 2003 Genetic diversity in yeast assessed with whole-genome oligonucleotide arrays. Genetics 163: 79–89.

Yin, J.-w., and G. Wang, 2014 The Mediator complex: a master coordinator of transcription and cell lineage development. Development 141: 977–987.

Young, E., H. Z. Abid, P.-Y. Kwok, H. Riethman and M. Xiao, 2020 Comprehensive Analysis of Human Subtelomeres by Whole Genome Mapping. PLOS Genetics 16: e1008347.

Yue, J. X., J. Li, L. Aigrain, J. Hallin, K. Persson et al., 2017 Contrasting evolutionary genome dynamics between domesticated and wild yeasts. Nat Genet 49: 913–924.

Zambrano, R., O. Conchillo-Sole, V. Iglesias, R. Illa, F. Rousseau et al., 2015 PrionW: a server to identify proteins containing glutamine/asparagine rich prion-like domains and their amyloid cores. Nucleic Acids Research 43: W331–W337.

Zhang, A., Z. Liu and L. C. Myers, 2013 Differential regulation of white-opaque switching by individual subunits of Candida albicans mediator. Eukaryotic Cell 12: 1293–1304.

Zhang, A., K. O. Petrov, E. R. Hyun, Z. Liu, S. A. Gerber et al., 2012 The Tlo proteins are stoichiometric components of Candida albicans Mediator anchored via the Med3 subunit. Eukaryotic Cell 11: 874–884.

Zhao, Z., and E. Boerwinkle, 2002 Neighboring-nucleotide effects on single nucleotide polymorphisms: a study of 2.6 million polymorphisms across the human genome. Genome research 12: 1679–1686.

Zhou, S. S., Z. Xing, H. Liu, X. G. Hu, Q. Gao et al., 2019 In-depth transcriptome characterization uncovers distinct gene family expansions for Cupressus gigantea important to this long-lived species’ adaptability to environmental cues. BMC Genomics 20: 1–14.

Zhu, X., L. Chen, J. O. Carlsten, Q. Liu, J. Yang et al., 2015 Mediator tail subunits can form amyloid-like aggregates in vivo and affect stress response in yeast. Nucleic Acids Res 43: 7306–7314.

Zhu, Y. O., M. L. Siegal, D. W. Hall and D. A. Petrov, 2014 Precise estimates of mutation rate and spectrum in yeast. Proceedings of the National Academy of Sciences 111: E2310–E2318.

